# Association between DNA methylation and ADHD symptoms from birth to school age: A prospective meta-analysis

**DOI:** 10.1101/806844

**Authors:** Alexander Neumann, Esther Walton, Silvia Alemany, Charlotte Cecil, Juan Ramon González, Dereje D. Jima, Jari Lahti, Samuli T. Tuominen, Edward D. Barker, Elisabeth Binder, Doretta Caramaschi, Ángel Carracedo, Darina Czamara, Jorunn Evandt, Janine F. Felix, Bernard F. Fuemmeler, Kristine B. Gutzkow, Cathrine Hoyo, Jordi Julvez, Eero Kajantie, Hannele Laivuori, Rachel Maguire, Léa Maitre, Susan K. Murphy, Mario Murcia, Pia M. Villa, Gemma Sharp, Jordi Sunyer, Katri Raikkönen, Marian Bakermans-Kranenburg, Marinus van IJzendoorn, Mònica Guxens, Caroline L. Relton, Henning Tiemeier

**Author notes:** CORRESPONDING AUTHOR: Henning Tiemeier, Erasmus University Medical Center Rotterdam, Department of Child and Adolescent Psychiatry/Psychology, Room KP-2822, PO-Box 2060, 3000 CB Rotterdam, The Netherlands. These authors contributed equally to this work (Alexander Neumann, Esther Walton, Silvia Alemany).

## Abstract

Attention-deficit and hyperactivity disorder (ADHD) is a common childhood disorder with a substantial genetic component. However, the extent to which epigenetic mechanisms play a role in the etiology of the disorder is not known. We performed epigenome-wide association studies (EWAS) within the Pregnancy And Childhood Epigenetics (PACE) Consortium to identify DNA methylation sites associated with ADHD symptoms at two methylation assessment periods: birth and school-age. We examined associations of DNA methylation in cord blood with repeatedly assessed ADHD symptoms (age range 4-15 years) in 2477 children from five cohorts and DNA methylation at school-age with concurrent ADHD symptoms (age 7-11 years) in 2374 children from ten cohorts. CpGs identified with nominal significance (p<0.05) in either of the EWAS were correlated between timepoints (ρ=0.30), suggesting overlap in associations, however, top signals were very different. At birth, we identified nine CpGs that were associated with later ADHD symptoms (P<1*10^−7^), including ERC2 and CREB5. Peripheral blood DNA methylation at one of these CpGs (cg01271805 located in the promotor region of ERC2, which regulates neurotransmitter release) was previously associated with brain methylation. Another (cg25520701) lies within the gene body of CREB5, which was associated with neurite outgrowth and an ADHD diagnosis in previous studies. In contrast, at school-age, no CpGs were associated with ADHD with P<1*10^−7^. In conclusion, we found evidence in this study that DNA methylation at birth is associated with ADHD. Future studies are needed to confirm the utility of methylation variation as biomarker and its involvement in causal pathways.

## Introduction

Attention-deficit and hyperactivity disorder (ADHD) is a common neurodevelopmental disorder characterized by impulsivity, excessive activity and attention problems. Symptoms often become apparent during school-age with a world-wide prevalence of 5-7.5%.^1^ Genetic heritability is estimated between 64%-88%.^2,3^ Additionally several environmental factors are suspected to impact ADHD, e.g. prenatal maternal smoking or lead exposure.^4–7^ However, the genetics and environmental pathways contributing to ADHD risk remain unclear. Possibly, DNA methylation, an epigenetic mechanism regulating gene expression, may mediate genetic or environmental effects.

Several studies have investigated DNA methylation in relation to ADHD diagnoses or symptoms using candidate approaches or epigenome-wide association studies (EWAS) in peripheral blood and saliva tissue.^8,9^ A leading hypothesis concerning the etiology of ADHD suggests that deficiencies in the dopamine system of the brain impact ADHD development.^4,10^ Consequently, candidate studies have focused on genes related to dopamine function. For instance, DNA methylation alterations in *DRD4*^*11–13*^, *DRD5*^*12*^, and *DAT1*^*12,14*^ genes have been associated with ADHD, though not consistently^15^. Beyond the candidate gene approach, three studies tested DNA methylation across the whole genome. One study performed an EWAS with saliva samples in school-aged children using a case-control design.^16^ The study identified differentially methylated probes in *VIPR2*, a gene expressed in the caudate and previously associated with psychopathology. Another EWAS investigated cord and peripheral blood DNA methylation at birth and at 7 years of age.^17^ At birth, 13 probes located in SKI, ZNF544, ST3GAL3 and PEX2 were associated with ADHD trajectories from age 7 to 15 years, but the methylation status of these probes at age 7 was not associated with ADHD cross-sectionally. An EWAS in adults with ADHD failed to find any differentially methylated sites in peripheral blood.^18^

Large multi-center epigenome-wide studies, which allow for increased power and generalizability, are lacking for childhood. Here we performed the first epigenome-wide prospective meta-analysis to identify DNA methylation sites associated with childhood ADHD symptoms in cohorts from the Pregnancy And Childhood Epigenetics (PACE) Consortium^19^. Since the temporal stability of methylation potentially associated with ADHD symptoms is unclear, we tested DNA methylation both at birth using cord blood and in school-age (age 7-9 years) using DNA derived from peripheral whole blood. In the analyses of cord blood methylation, the aim was to explain ADHD symptoms between ages 4 and 15 years. Many participating cohorts assessed ADHD repeatedly and we employed a repeated measures design to increase precision. Furthermore, we utilized data in childhood to examine cross-sectional DNA methylation patterns associated with ADHD symptoms at school age.

## Materials and methods

This study comprises a birth methylation EWAS and a school-age methylation EWAS described successively below.

### Birth Methylation EWAS

#### Participants

Five cohorts (Avon Longitudinal Study of Parents and Children (ALSPAC),^20–22^ Generation R (GENR),^23^ INfancia y Medio Ambiente (INMA),^24^ Newborn Epigenetic Study (NEST)^25,26^ and Prediction and prevention of preeclampsia and intrauterine growth restriction (PREDO)^27^) in the PACE consortium had information on DNA methylation in cord blood and ADHD symptoms. These cohorts have a combined sample size of 2477 (Table 1). Participants were mostly of European ancestry, except for NEST, an American cohort which also included participants of African ancestry. In NEST, separate EWAS were conducted for participants identifying as black or white to account for ancestry heterogeneity. See Supplementary Information 1 for full cohort descriptions.

**Table 1:** Cohort characteristics

|  |  |  |  |  |  | Standardized regression coefficients |  |  | BACON estimates |  |  |
| --- | --- | --- | --- | --- | --- | --- | --- | --- | --- | --- | --- |
| Cohort | Ancestry/<br>Ethnicity | n | Methylation<br>Age | ADHD<br>Age | Instrument<br>(Age) | 33% | 50% | 66% | $\lambda$ | Inflation | Bias |
| *Birth EWAS* |  |  |  |  |  |  |  |  |  |  |  |
| ALSPAC | European | 714 | 0 | 8, 11, 14, 15 | DAWBA | -0.21 | 0.25 | 0.89 | 1.60 | 1.10 | 0.37 |
| GENR | European | 1191 | 0 | 6,8,10 | CBCL (6,10),<br>Conners (8) | -0.48 | 0.01 | 0.53 | 1.51 | 1.20 | 0.05 |
| INMA | European | 325 | 0 | 7,9 | Conners (7),<br>CBCL (9) | -1.37 | -0.40 | 0.43 | 0.80 | 0.87 | -0.19 |
| NEST | Black | 55 | 0 | 5 | BASC | -3.50 | -0.03 | 3.63 | 1.16 | 1.10 | 0.00 |
| NEST | White | 56 | 0 | 5 | BASC | -2.54 | -0.09 | 2.36 | 0.80 | 0.92 | -0.01 |
| PREDO | European | 136 | 0 | 5 | Conners | -1.55 | -0.25 | 1.20 | 1.45 | 0.95 | 0.21 |
| META | - | 2477 | - | - | - | -0.37 | 0.02 | 0.42 | 1.86 | 1.10 | 0.01 |
| *School-age EWAS* |  |  |  |  |  |  |  |  |  |  |  |
| ALSPAC | European | 651 | 7 | 8 | DAWBA | -0.61 | -0.10 | 0.54 | 1.09 | 1.00 | -0.08 |
| GENR | European | 395 | 10 | 10 | CBCL | -0.93 | -0.00 | 0.98 | 1.00 | 0.97 | -0.01 |
| GLAKU | European | 215 | 12 | 12 | CBCL | -0.79 | 0.31 | 1.50 | 0.92 | 0.96 | 0.13 |
| HELIX | European | 1034 | 8 | 8 | CBCL | -0.26 | 0.47 | 1.40 | 1.11 | 0.98 | 0.28 |
| HELIX | Pakistani | 79 | 7 | 7 | CBCL | -1.66 | 1.86 | 5.48 | 0.98 | 0.96 | 0.26 |
| Meta | - | 2374 | - | - | - | -0.24 | 0.14 | 0.62 | 0.96 | 0.92 | 0.14 |

**n** Number of participants
**33%, 50%, 66%** Quartiles of regression coefficient distribution
**$\lambda$** Inflation of p-values
**Inflation** Inflation of p-values due to suspected bias
**Bias** Trend toward negative/positive distribution of regression coefficients due to suspected bias

#### DNA Methylation and QC

DNA methylation in cord blood was measured using the Illumina Infinium HumanMethylation450K BeadChip (Table S1). Methylation levels outside of the lower quartile minus 3*interquartile or upper quartile plus 3*interquartile range were removed. Each cohort ran the EWAS separately according to a pre-specified harmonized analysis plan. The distribution of the regression estimates and p-values were examined for each cohort and pooled results. Deviations from a normal distribution of regression estimates or a higher number of low p-values than expected by chance may be signs of residual confounding, or the result of a true poly-epigenetic signal. To help in interpretation of the results, we used the BACON method.^28^ BACON analyzes the distribution of regression coefficients and estimates an empirical null distribution. Results can then be compared against the empirical null, which already includes biases, rather than the theoretical null. We excluded CpG probes, that were available in fewer than four cohorts, fewer than 1000 participants, and allosomal probes, due to the complex interpretation of dosage compensation.

#### ADHD Symptoms

ADHD symptoms were measured when children were 4-15 years old (depending on the cohort) with parent-rated instruments, specifically the Behavior Assessment System for Children (BASC),^29^ Child Behavior Checklist (CBCL),^30,31^ Conners^32^ and the Development and Well-Being Assessment (DAWBA)^33^ (Table S2). If a cohort had measured ADHD symptoms repeatedly (3 cohorts), we used a mixed model (see statistical analysis). The repeated measure design increased the precision of the ADHD severity estimate and sample size, since missing data in an assessment can be handled with maximum likelihood. Given the variety of instruments used within and across cohorts, all ADHD scores were z-score standardized to enable meta-analysis.

#### Statistical analysis

Cohorts with repeated ADHD assessment were analyzed using linear mixed models, with z-scores of ADHD symptoms as the outcome and methylation (in betas, ranging from 0 (unmethylated) to 1 (methylated)) as the main predictor. Each CpG probe was analyzed separately and pooled p-values were adjusted for multiple correction using Bonferroni adjustment. We used a random intercept on the participant and batch level, to account for clustering due to repeated measures and batch effects. The following potential confounders were included as fixed effects: maternal age, educational level, smoking status (yes vs no during pregnancy), gestational age, sex, and estimated white blood cell proportions (Bakulski reference estimated with the Houseman method).^34^ Mixed models were fitted using restricted maximum likelihood. We used R^35^ with the lme4^36^ package to estimate the models. Cohorts with a single ADHD assessment wave used a model without random effects or batch level only.

Meta-analysis was performed using the Han and Eskin random effects model.^37^ This model does not assume that true effects are homogeneous between cohorts, however, it does assume that null effects are homogeneous. This modified version of the random effect model has comparable power to a fixed effects analysis, while better accounting for study heterogeneity, such as ancestry differences, in simulation studies.^37^ Genome-wide significance was defined at the Bonferroni-adjustment threshold of p<1*10^−7^, suggestive significance at p<1*10^−5^, and nominal significance at p<0.05.

#### Follow-up analyses

We performed several look-ups of genome-wide significant probes. We used the BECon database^38^ to check the correlation between peripheral and brain methylation levels in post-mortem tissue. To test genetic influence we interrogated the genome-wide significant probes in MeQTL^39^ and twin heritability databases.^40^ We also attempted to replicate genome-wide significant probes reported in a previous EWAS from the ALSPAC study.^17^ For replication we reran the meta-analysis without the ALSPAC cohort. To quantify the variance explained by genome-wide significant probes, we predicted ADHD scores at age 8 in Generation R by all meta-analytically genome-wide significant probes. We applied 10-fold cross-validation with 100 repetitions to improve generalizability and reduce bias from Generation R, which was part of the discovery.

#### Pathway Analysis

Pathway enrichment analysis were performed with the missMethylpackage^41^ on suggestive probes (P<1*10^−5^). We used as references: gene ontology (GO), KEGG and curated gene sets (http://software.broadinstitute.org/gsea/msigdb/collections.jsp#C2) from the Broad Institute Molecular signatures database^42^. P-values were adjusted using the default procedures by the number of CpGs associated with each gene^43^ and false discovery rate.

To test enrichment for regulatory features (gene relative position, CpG island relative position and blood chromatin states) we applied χ^2^ tests. Enrichment tests were performed for all CpGs, hypo and hypermethylated CpGs separately. CpG annotation was performed with the IlluminaHumanMethylation450kanno.ilmn-12.hg19 R package.^44^ Annotation to chromatin states was from the Roadmap Epigenomics Project (https://egg2.wustl.edu/roadmap/web_portal/). See Supplementary Information 2 for full description.

### School-age methylation EWAS

#### Participants

Nine cohorts (ALSPAC, GENR, HELIX^45^ and GLAKU^46^) with a combined sample size of 2374 joined the school-age methylation EWAS (Table 1, Supplementary Information 1). HELIX consists of six jointly analyzed sub-cohorts^45^ All cohorts had participants of European ancestry, except HELIX, which also included participants with a Pakistani background living in the UK, which were treated as a separate cohort in the meta-analysis. Fifty-three percent of participants in the school-age EWAS were also part of the birth EWAS.

#### DNA Methylation and QC

DNA methylation was measured at ages 7-12 in peripheral whole blood. The Illumina Infinium HumanMethylation450K BeadChip and Infinium MethylationEPIC Kit (GLAKU) were used to interrogate CpG probes. QC steps were identical to the birth methylation EWAS.

#### ADHD Symptoms

ADHD symptoms were measured at the same age as DNA methylation (age 7-11 years) with the parent-rated measures DAWBA and CBCL (Table S2). Only the assessment closest to the DNA methylation assessment age was analyzed.

#### Statistical analysis

The statistical model was similar to the model used in the birth methylation EWAS without participant level random effect. However, cell counts were estimated with the Houseman method using the Reinius reference.^47^. We also added assessment age as covariate. The meta-analysis methods were identical to the birth methylation EWAS.

#### Follow-up analyses

We did not perform follow-up analyses due low signal. However, we attempted to replicate six probes identified as suggestive in a previous case-control EWAS in school-age.^16^

## Results

### Birth Cord Blood Methylation

#### EWAS Quality Check

Four out of the six cohorts showed larger number of low p-values than expected under the null, as indexed by high λ (Table 1). BACON analysis suggested that the majority of the inflation was due to a true signal, as indicated by inflation values clearly lower than λ. To test the impact of sample size on λ, we restricted the GENR sample randomly to 900 and 1100 participants, resulting in 812 and 991 participants due to missing covariates. The lambdas were 0.96, 1.21, 1.51 for 812, 991, and 1191 participants. We thus conclude that the over-representation of low p-values is mostly due to sufficient power to detect associations at higher sample sizes.

The BACON analyses also indicated a trend towards positive/negative regression coefficients in some of the datasets, which might indicate confounding, e.g. by population stratification. To test this, we added principal components of ancestry in GENR and ALSPAC, but these did not meaningfully change results.

We conducted the meta-analysis under the assumption that any such biases will be corrected in the pooled analysis, since they were not homogeneous across cohorts. Indeed, the pooled estimates did not show a trend towards positive or negative regression estimates (Median=+0.02), only an overrepresentation of low p-values (λ=1.86, Figure 1). The BACON estimates for inflation suggested that these are mostly due to a true signal (Inflation=1.1).

**Figure 1:**
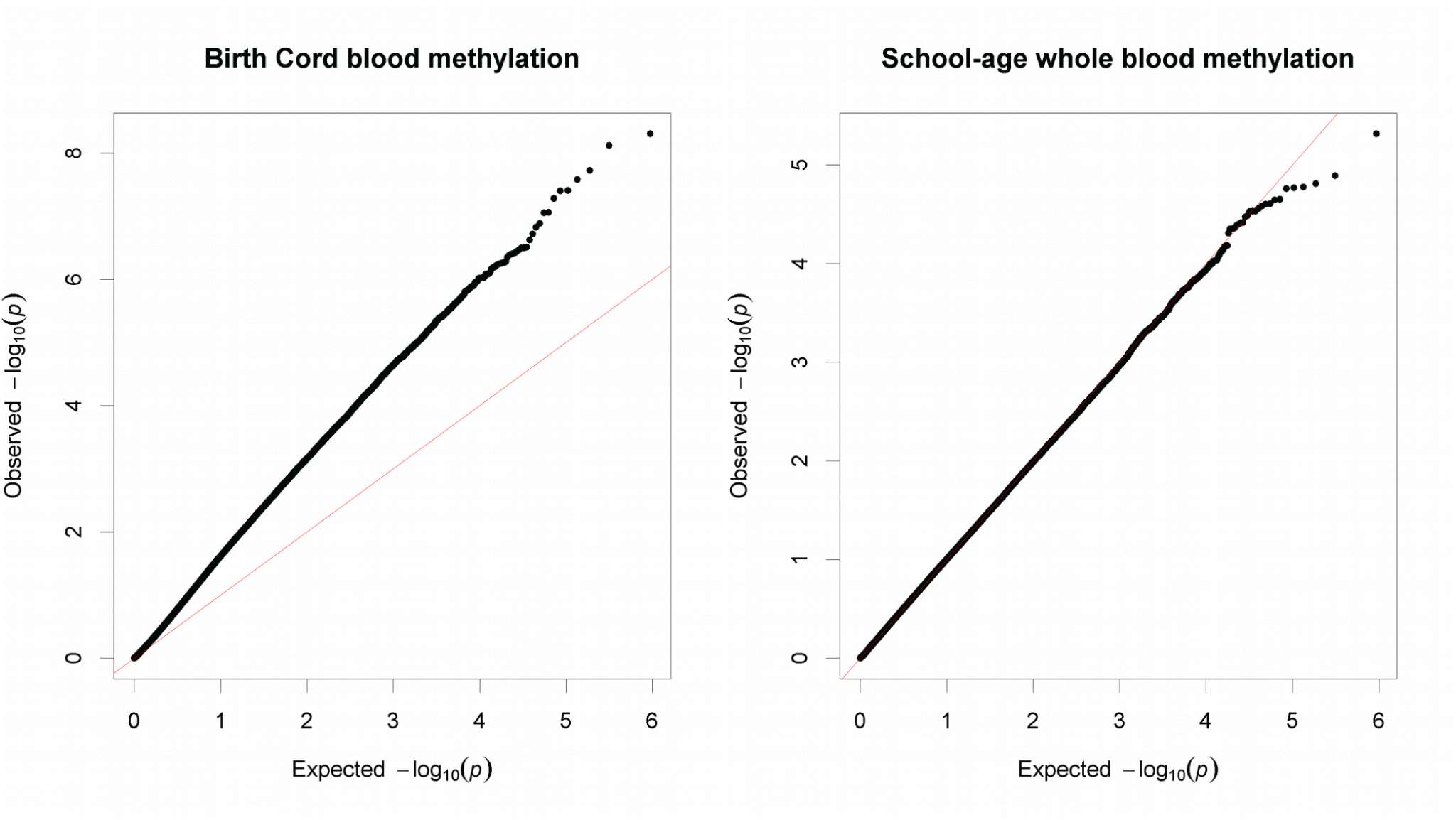
Quantile-quantile plot of observed −log_10_ p-values in the cord blood and school-age EWAS vs expected −log_10_ p-values under assumption of chance findings only. The diagonal line represents the distribution of the expected p-values under the null. Points above the diagonal indicate p-values which are lower than expected.

#### Single Probe Analysis

After QC, 472,817 CpG sites remained for the meta-analysis. Results of the cord blood EWAS are shown in Figure 2. Nine CpG sites showed genome-wide significance (p<1*10^−7^, Table 2). ADHD symptoms were between 0.16SD (SE=0.03) and 0.44SD (SE= 0.12) higher with 10% lower methylation at these probes. Eight probes out of nine that were available in the BECon database^38^ are typically methylated in both whole blood and the brain (Figures 3, S1 and S2). A lookup in the BECon database revealed that the CpG site cg01271805 in the promoter region of gene *ERC2* shows variable methylation in three brain regions (BA10, BA20, BA7). Importantly, methylation levels in the brain are moderately correlated with whole blood methylation (ρ=0.33-0.46) (Figure 3), suggesting that peripheral cg01271805 methylation levels are a useful marker for brain methylation levels. The other seven genome-wide significant probes showed less consistent correlations between blood and brain tissues and associated genes had less specificity for expression in the brain, based on GTEx^48^ data. No SNP was associated with our nine top CpG probes when accounting for linkage disequilibrium according to the MeQTL database^39^. Furthermore, all nine probes had a twin heritability below 20% in a previous study (Table S3).^40^ After adjusting for inflation and bias with BACON, only one CpG remained statistically significant (cg25520701, CREB5, ß =-3.54, SE = 0.66, p = 9.59*10^−8^). It should be noted, that the BACON adjusted p-values rely on statistics from the traditional random effects model. With the traditional model, only cg25520701, cg09762907 and cg22997238 remained genome-wide significant. Thus the difference in p-value is not solely the result of adjustment for the inflation, but also the use of more conservative tests. In Generation R, the joint explained variance of ADHD scores at age 8 by the genome-wide significant probes was 1.8% (R^2^ from 10-fold repeated cross-validation).

**Table 2:** EWAS Results

| CpG | Gene | Chr | Position | Birth methylation |  |  |  |  | School-age methylation |  |  |  |  |
| --- | --- | --- | --- | --- | --- | --- | --- | --- | --- | --- | --- | --- | --- |
|  |  |  |  | n<sub>studies</sub> | n | B | SE | p | n<sub>studies</sub> | n | B | SE | p |
| cg25520701 | CREB5 | 7 | 28800657 | 6 | 2450 | -3.53 | 0.60 | 4.95E-09 | 5 | 2279 | -0.13 | 1.09 | 0.94 |
| cg24838839 | Intergenic | 5 | 61031569 | 6 | 2468 | -4.15 | 1.79 | 3.95E-08 | 5 | 2287 | 1.52 | 1.38 | 0.33 |
| cg22997238 | Intergenic | 7 | 36014218 | 6 | 2465 | -1.63 | 0.30 | 8.81E-08 | 5 | 2291 | -0.06 | 0.47 | 0.94 |
| cg21600027 | Intergenic | 4 | 124443502 | 6 | 2464 | -3.04 | 0.81 | 2.64E-08 | 5 | 2281 | 0.98 | 0.89 | 0.33 |
| cg17876201 | ZBTB38 | 3 | 141139991 | 6 | 2457 | -4.41 | 1.20 | 7.58E-09 | 4 | 2066 | 0.56 | 1.32 | 0.73 |
| cg11251614 | PPIL1 | 6 | 36839846 | 6 | 2451 | -3.43 | 0.68 | 3.89E-08 | 5 | 2276 | 0.77 | 1.52 | 0.68 |
| cg09762907 | TRERF1 | 6 | 42290256 | 6 | 2460 | -2.11 | 0.39 | 8.76E-08 | 5 | 2284 | -0.55 | 0.64 | 0.46 |
| cg09158638 | Intergenic | 16 | 62309996 | 6 | 2470 | -2.55 | 1.40 | 1.89E-08 | 5 | 2270 | -0.33 | 1.04 | 0.80 |
| cg01271805 | ERC2 | 3 | 55694954 | 6 | 2469 | -2.86 | 1.71 | 5.24E-08 | 5 | 2289 | 0.28 | 0.73 | 0.76 |

**Chr** Chromosome**n<sub>studies</sub>** Number of studies**n** Number of participants**B** Regression coefficient**SE** Standard error

**Figure 2:**
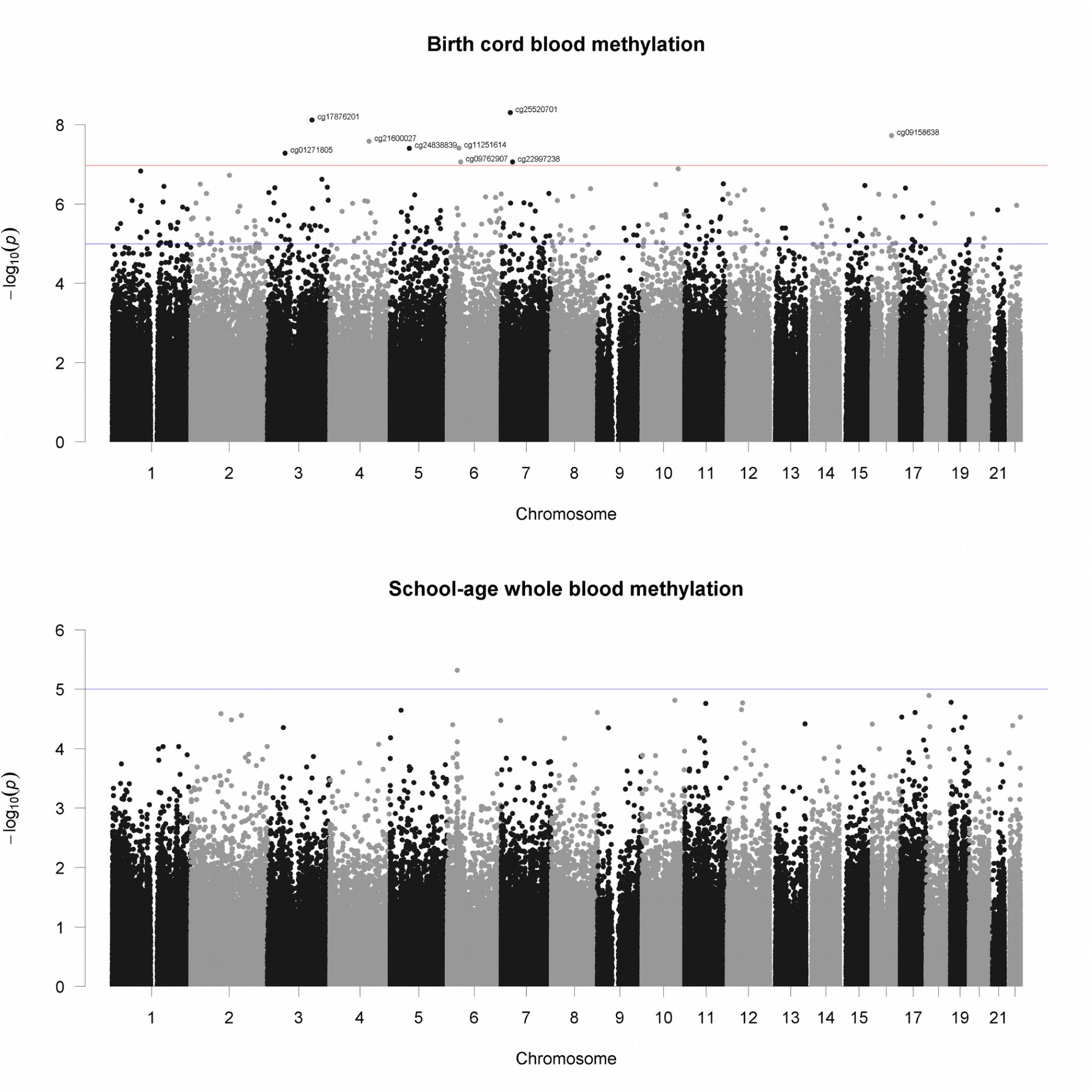
Manhattan plot of −log_10_ p-values vs CpG position (basepair and chromosome). Red line indicates genome-wide significant (p<1*10^−7^) and blue line suggestive threshold (p<1*10^−5^).

**Figure 3:**
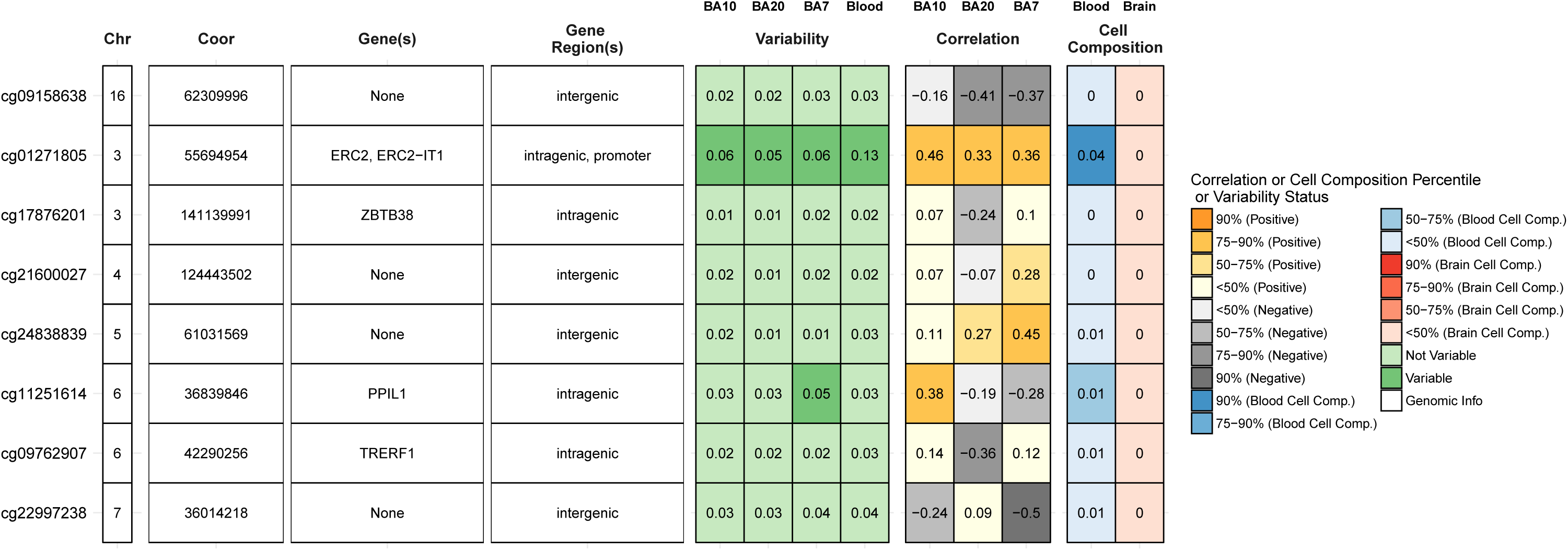
Lookup of brain-blood correlations and variability of genome-wide significant CpG sites in the BECon database.

#### Pathway Analysis

Two-hundred forty-nine probes showed suggestive (P<1*10^−5^) associations and were annotated to 182 unique genes. In gene-based analyses no pathway survived multiple testing correction.

The 248 suggestive CpGs were enriched in intergenic regions. Of these, hypomethylated CpGs were enriched for 3’UTR regions and depleted for TSS200 and first exon regions, open sea, north shelf and south shelf regions, south shore and islands. Regarding chromatin states, hypomethylated probes showed an enrichment for transcription (Tx and TxWk), quiescent positions and depletion for transcription start site positions (TSSA, TxFlnk, TxFlnk), bivalent (EnhBiv) and repressor (ReprPC) positions. Hypermethylated probes showed the opposite enrichment/depletion patterns. See Supplementary Information 2 for full results.

#### Replication of previous EWAS

We attempted to replicate findings for 13 CpGs, at which DNA methylation at birth was associated with ADHD trajectories.^17^ However, no probe survived multiple-testing correction. (Table S4).

### School-age methylation

#### EWAS Quality Checks

The regression coefficient distribution showed no signs of errors, but three out of the five cohorts showed a trend towards positive associations in separate analyses (Table 1). The lambda was below 1.11 for all cohorts. BACON suggested no inflation of the test statistics due to confounding or other biases, though the trend towards positive associations remained. The pooled results showed a low lambda (λ=0.96), no inflation (BACON inflation estimate = 0.92), but a slight over-representation of positive associations (BACON bias estimate = 0.14).

#### Single Probe Meta-Analysis

We associated DNA methylation at school-age in whole-blood at 466,574 CpG sites with ADHD symptoms at the same age. No CpG reached genome-wide significance (all p>4.96E-06, Figure 2). Furthermore, none of the loci at which DNA methylation at birth was significantly associated with ADHD symptoms, also showed a cross-sectional association at school-age (p>0.33).

#### Replication of previous EWAS

We attempted to replicate the six most suggestive EWAS CpGs of a previous case-control study.^16^ While all but one showed a consistent direction, none of the CpGs were statistically significant. (Table S5)

#### Stability of methylation association across age

The associations between methylation at birth with ADHD symptoms and methylation at school-age with ADHD symptoms were largely consistent for nominally significant probes. The regression estimates from CpG sites, with nominally significant associations at birth (p<0.05, n=73,057) correlated with the regression estimates of the school-age EWAS (ρ=0.45). When restricting the school-age methylation EWAS to those cohorts, which were not featured in the birth methylation EWAS (thus excluding overlaps), the correlation remained (ρ=0.30). Vice versa, when filtering for probes which were nominally significant at school-age, 23,770 probes remained of which 4075 overlapped with nominally significant probes at birth. The correlation for this set was very similar, ρ=0.47 among all cohorts and ρ=0.35 between independent cohorts.

## Discussion

In this population-based study, we performed the first epigenome-wide meta-analysis of ADHD symptoms in childhood, using two DNA methylation assessments (birth and school-age), as well as repeated measures of ADHD symptoms. DNA methylation at birth, but not at school-age, was associated with later development of ADHD symptoms with genome-wide significance at nine loci. Interestingly, the identified probes showed a pattern of a high average rate of methylation in cord blood, while lower levels of methylation were associated with more ADHD symptoms in childhood. DNA methylation in cord blood reflects the effects of genetics and the intrauterine environment. The results suggest that cord blood DNA methylation is a marker for some of the ADHD risk factors before birth or functions as a potential mediator of these risk factors. While not impossible, reverse causality at this age is unlikely to explain our results, as ADHD only manifests at later stages of development.

We analyzed DNA methylation in cord and peripheral blood, which may not correspond to the methylation status in the brain. DNA methylation in the brain arguably has the strongest a priori likelihood of representing causal mechanisms. Seven out of eight significant probes did not show consistent correlation between methylation status in whole blood and post-mortem brain tissue in a previous study, i.e. DNA methylation levels in blood may not represent brain levels and thus associations with ADHD may be different.^38^ However, methylation levels of cg01271805 in whole blood are associated with methylation levels in various brain regions. Importantly, this probe lies in the promoter region of the gene *ERC2*, that is highly expressed in brain tissue. *ERC2* regulates calcium dependent neurotransmitter release in the axonal terminal.^49^ Specifically, *ERC2* is suspected to increase the sensitivity of voltage dependent calcium channels to hyperpolarization, resulting in higher neurotransmitter release. SNPs in the *ERC2* locus have been suggested to distinguish schizophrenia and bipolar disorder patients^50^ and to impact cognitive functioning^51^. *ERC2* is especially expressed in Broadmann area 9 of the frontal cortex.^48^ Previous imaging studies have demonstrated differential activation in this area when children with or without ADHD performed various cognitive tasks.^52,53^ The correlation with brain methylation, the location in a promoter and gene expression in the brain make cg01271805 a plausible candidate locus, where reduced methylation may be mechanistically involved in ADHD development. We hypothesize, that lower methylation levels at cg01271805 increases the expression of *ERC2*, which in turn increases neurotransmitter release, with an adverse impact on the development of ADHD symptoms. Another gene with a genome-wide significant probe and high relevance for neural functioning is *CREB5* (cg25520701). *CREB5* is expressed in fetal brain and the prefrontal cortex, and has been previously related to neurite outgrowth. Moreover, SNPs in this gene were associated with ADHD in two recent GWAS.^54,55^ Thus, it is plausible that differences in DNA methylation at this locus may modify ADHD risk during development.

While the birth methylation EWAS identified several loci, associating school-age methylation with concurrent ADHD symptoms revealed no genome-wide significant associations. Furthermore, the overall association signal was lower, despite similar sample sizes. None of the probes, which were significantly associated at birth showed any association when measured at school-age. Given that sample sizes were comparable, this difference must come from changes in the epigenome or study heterogeneity, rather than differences in statistical power. In terms of instrument heterogeneity, the school-age EWAS was more homogeneous, almost exclusively using CBCL. Additionally, as both EWAS feature a mix of several cohorts selected based on the same criteria and around half of the participants were represented at both time points, study heterogeneity appears to be an unlikely explanation. The stronger signal in the birth EWAS may be considered surprising given that typically two measures are typically more strongly associated if measured in closer temporal proximity. However, in line with our results Walton et al. also observed in a previous EWAS,^17^ that birth methylation may be a better predictor of later ADHD symptoms than childhood methylation, possibly reflecting sensitive periods. Whether DNA methylation in cord blood has stronger causal effects or is a better marker for early life factors cannot be concluded from the present study. Alternatively, tissue differences between cord blood and whole blood may account for the differences in association pattern. Finally, it is possible that interventions in childhood and other environmental influences reduced the initial epigenetic differences at birth between children with higher and lower ADHD symptoms. Yet, we observed consistency in the associations of methylation at both timepoints with ADHD symptoms. The regression estimates of both EWAS correlated on a genome-wide level.

Strengths of this study include the large sample size, repeated outcome measures, Spanish Institute of Health Carlos III extensive control for potential confounders and the use of DNA methylation at two different time-points, enabling us to characterize both prospective and cross-sectional associations with ADHD symptoms. However, several limitations need to be discussed as well. A causal interpretation of our findings is challenged by the possibility of residual confounding and reverse causality. DNA methylation might be a marker for untested adverse environmental factors that could affect ADHD via independent pathways. In addition, children with higher ADHD symptoms may evoke a particular environment, which might shape the epigenome. Larger sample sizes are necessary to detect further methylation sites. As is typical for (epi-)genetic studies, the effect size of individual top probes was rather small: the joint effect of the genome-wide probes was estimated below 2%. However, the strong genome-wide epigenetic signal suggests a potential for the development of epigenetic-scores based on birth methylation, which could lead to early prevention efforts before ADHD symptoms arise.

In summary, we identified nine CpG sites for which lower methylation status at birth is associated with later development of ADHD symptoms. The results suggest that DNA methylation in *ERC2* and *CREB5* may exert an influence on ADHD symptoms, potentially via modification of neurotransmitter functioning or neurite outgrowth.

## Acknowledgments

We thank all the children and families who took part in this study, as well as the support of hospitals, midwives and pharmacies.

## ALSPAC

We are grateful to the whole ALSPAC team, which includes interviewers, computer and laboratory technicians, clerical workers, research scientists, volunteers, managers, receptionists and nurses. The UK Medical Research Council (MRC) and Wellcome (Grant ref: 102215/2/13/2) and the University of Bristol provide core support for ALSPAC. This publication is the work of the authors and E Walton will serve as guarantors for the contents of this paper. A comprehensive list of grants funding is available on the ALSPAC website (http://www.bristol.ac.uk/alspac/external/documents/grant-acknowledgements.pdf). Methylation data in the ALSPAC cohort were generated as part of the UK BBSRC funded (BB/I025751/1 and BB/I025263/1) Accessible Resource for Integrated Epigenomic Studies (ARIES, http://www.ariesepigenomics.org.uk). Edward Barker is supported by an ESRC grant ES/S013229/1. Doretta Caramaschi, Gemma Sharp and Caroline L. Relton work in a department supported by the UK MRC (MC_UU_00011/5). Gemma Sharp is further supported by an MRC New Investigator Research Grant (MR/S009310/1) and the European Joint Programming Initiative “A Healthy Diet for a Healthy Life” (JPI HDHL, NutriPROGRAM project, MRC, MR/S036520/1).

## GENR

The Generation R Study is conducted by the Erasmus Medical Center in close collaboration with the Erasmus University Rotterdam, Faculty of Social Sciences, the Municipal Health Service Rotterdam area, the Rotterdam Homecare Foundation, Rotterdam and the Stichting Trombosedienst & Artsenlaboratorium Rijnmond (STAR-MDC), Rotterdam. We gratefully acknowledge the contribution of general practitioners, hospitals, midwives and pharmacies in Rotterdam. The generation and management of the Illumina 450K methylation array data (EWAS data) for the Generation R Study was executed by the Human Genotyping Facility of the Genetic Laboratory of the Department of Internal Medicine, Erasmus MC, the Netherlands. We thank Mr. Michael Verbiest, Ms. Mila Jhamai, Ms. Sarah Higgins, Mr. Marijn Verkerk and Dr. Lisette Stolk for their help in creating the EWAS database. We thank Dr. A.Teumer for his work on the quality control and normalization scripts.The general design of the Generation R Study is made possible by financial support from Erasmus Medical Center, Rotterdam, Erasmus University Rotterdam, the Netherlands Organization for Health Research and Development (ZonMw) and the Ministry of Health, Welfare and Sport. The EWAS data was funded by a grant from the Netherlands Genomics Initiative (NGI)/Netherlands Organisation for Scientific Research (NWO) Netherlands Consortium for Healthy Aging (NCHA; project nr. 050-060-810), by funds from the Genetic Laboratory of the Department of Internal Medicine, Erasmus MC, and by a grant from the National Institute of Child and Human Development (R01HD068437). A. Neumann and H. Tiemeier are supported by a grant of the Dutch Ministry of Education, Culture, and Science and the Netherlands Organization for Scientific Research (NWO grant No. 024.001.003, Consortium on Individual Development). A. Neumann is also supported by a Canadian Institutes of Health Research team grant. The work of H. Tiemeier is further supported by a NWO-VICI grant (NWO-ZonMW: 016.VICI.170.200). M.H. van IJzendoorn and M.J. Bakermans-Kranenburg were supported by the Netherlands Organization for Scientific Research (SPINOZA, VICI), and M.J. Bakermans-Kranenburg was supported by the European Research Council (AdG 669249). J.F.F. has received funding from the European Joint Programming Initiative “A Healthy Diet for a Healthy Life” (JPI HDHL, NutriPROGRAM project, ZonMw the Netherlands no.529051022). This project received funding from the European Union’s Horizon 2020 research and innovation programme (733206, LifeCycle; 633595, DynaHEALTH.

## GLAKU

We thank all the research nurses, research assistants, and laboratory personnel involved in the GLAKU study. The study has been supported by Academy of Finland, University of Helsinki, Hope and Optimism Initiative, Finnish Foundation for Pediatric Research, Sigrid Juselius Foundation, Jalmari and Rauha Ahokas Foundation, Signe and Ane Gyllenberg Foundation, Yrjo Jahnsson Foundation, Juho Vainio Foundation, Emil Aaltonen Foundation, and Ministry of Education and Culture, Finland.

## INMA

A full roster of the INMA Project Investigators can be found at http://www.proyectoinma.org/presentacion-inma/listado-investigadores/en_listado-investigadores.html. Silvia Alemany is funded by a Juan de la Cierva-Incorporación fellowship (IJCI-2017-34068) awarded by the Spanish Ministerio de Economía, Industria y Competitividad. Mònica Guxens is funded by a Miguel Servet fellowship (MS13/00054, CP18/00018) awarded by the Spanish Institute of Health Carlos III. ISGlobal is a member of the CERCA Programme, Generalitat de Catalunya. Jordi Julvez is funded by a Miguel Servet fellowship (MS14/00108, CP14/00108) awarded by the Spanish Institute of Health Carlos III.

## HELIX

The authors would like to thank allpractitioners and researchers in the six countries who took part in this study. The authors would like to thank Sonia Brishoual, Angelique Serre and Michele Grosdenier (Poitiers Biobank, CRB BB-0033-00068, Poitiers, France) for biological sample management and Professor Frederic Millot (Principal Investigator), Elodie Migault, Manuela Boue and Sandy Bertin (Clinical Investigation Center, Inserm CIC1402, CHU de Poitiers, Poitiers, France) for planning and investigational actions. The authors would like to thank Veronique Ferrand-Rigalleau, Céline Leger and Noella Gorry (CHU de Poitiers, Poitiers, France) for administrative assistance (EDEN). The authors would like to thank Silvia Fochs, Nuria Pey, Cecilia Persavente and Susana Gross for field work, sample management and overall management in INMA. The authors would like to thank Georgia Chalkiadaki and Danai Feida for biological sample management, to Eirini Michalaki, Mariza Kampouri, Anny Kyriklaki and Minas Iakovidis for field study performance and to Maria Fasoulaki for administrative assistance (RHEA). The authors would also like to thank Ingvild Essén for thorough field work, Heidi Marie Nordheim for biological sample management and the Norwegian Mother, Father and Child cohort study (MoBa) administrative unit. The MoBa Cohort Study is supported by the Norwegian Ministry of Health and Care Services and the Ministry of Education and Research. The contribution of the Spanish National Genotyping Center (CEGEN-PRB2) is also acknowledged.

## NEST

The NEST cohort has been supported by the National Institute of Environmental Health Sciences (R01ES016772 [CH], R21ES014947 [CH], P30ES025128 [CH, DDJ, RM], R01MD011746 [CH, SKM], P30ES011961 pilot project [SKM], P01ES022831 [SKM, BFF]), the US Environmental Protection Agency (RD-83543701 [SKM, BFF]), the National Institute of Diabetes and Digestive and Kidney Diseases (R01DK085173 [CH, SKM]), the National Institute of Aging (R21AG041048 [BFF]) and the Duke Cancer Institute [CH, SKM]. The contents are solely the responsibility of the authors and do not necessarily represent the official views of the National Institutes of Health or the United States Environmental Protection Agency (US EPA). Further, USEPA does not endorse the purchase of any commercial products or services mentioned in the publication.

## PREDO

The PREDO Study has been funded by the Academy of Finland, EraNet Neuron, EVO (a special state subsidy for health science research), University of Helsinki Research Funds, the Signe and Ane Gyllenberg foundation, the Emil Aaltonen Foundation, the Finnish Medical Foundation, the Jane and Aatos Erkko Foundation, the Novo Nordisk Foundation, the Päivikki and Sakari Sohlberg Foundation, the Sigrid Juselius Foundation granted to members of the Predo study board. Methylation assays were funded by the Academy of Finland. The PREDO study would not have been possible without the dedicated contribution of the PREDO study group members: A-K Pesonen (Department of Psychology and Logopedics, University of Helsinki; Finland), A Aitokallio-Tallberg, A-M Henry, VK Hiilesmaa, T Karipohja, R Meri, S Sainio, T Saisto, S Suomalainen-Konig, V-M Ulander, T Vaitilo (Department of Obstetrics and Gynaecology, University of Helsinki and Helsinki University Central Hospital, Helsinki, Finland), L Keski-Nisula, Maija-Riitta Orden (Kuopio University Hospital, Kuopio Finland), E Koistinen, T Walle, R Solja (Northern Karelia Central Hospital, Joensuu, Finland), M Kurkinen (Päijät-Häme Central Hospital, Lahti, Finland), P.Taipale. P Staven (Iisalmi Hospital, Iisalmi, Finland), J Uotila (Tampere University Hospital, Tampere, Finland). We also thank all the research nurses, research assistants, and laboratory personnel involved in the PREDO study.

