## Supplementary Information 1 for "Association between DNA methylation and ADHD symptoms from birth to school age: A prospective meta-analysis"

### Cohort descriptions

#### ALSPAC

Pregnant women residing in the study area of former county Avon, United Kingdom, with an expected delivery date between April 1991 and December 1992 were invited to enroll in the ALSPAC study. The initial number of pregnancies enrolled is 14,541. Of these initial pregnancies, there was a total of 14,676 foetuses, resulting in 14,062 live births and 13,988 children who were alive at 1 year of age. For more information on the study design has been published previously (Fraser et al. 2013, Boyd et al. 2013). Blood from 1018 mother–child pairs (children at three time points and their mothers at two time points) were selected for analysis as part of the Accessible Resource for Integrative Epigenomic Studies (ARIES, http://www.ariesepigenomics.org.uk/; Relton et al. 2015). The ALSPAC study website contains details of all the data that are available through a fully searchable data dictionary and variable search tool (http://www.bristol.ac.uk/alspac/researchers/our-data/). Ethical approval for the study was obtained from the ALSPAC Ethics and Law Committee and the Local Research Ethics Committees. Consent for biological samples has been collected in accordance with the Human Tissue Act (2004). Informed consent for the use of data collected via questionnaires and clinics was obtained from participants following the recommendations of the ALSPAC Ethics and Law Committee at the time

http://www.bristol.ac.uk/alspac/

Boyd A, Golding J, Macleod J, Lawlor DA, Fraser A, Henderson J, Molloy L, Ness A, Ring S, Davey Smith G. Cohort Profile: The ‘Children of the 90s’; the index offspring of The Avon Longitudinal Study of Parents and Children (ALSPAC). International Journal of Epidemiology 2013; 42: 111-127.

Fraser A, Macdonald-Wallis C, Tilling K, Boyd A, Golding J, Davey Smith G, Henderson J, Macleod J, Molloy L, Ness A, Ring S, Nelson SM, Lawlor DA. Cohort Profile: The Avon Longitudinal Study of Parents and Children: ALSPAC mothers cohort. International Journal of Epidemiology 2013; 42:97-110.

Caroline L Relton, Tom Gaunt, Wendy McArdle, Karen Ho, Aparna Duggirala, Hashem Shihab, Geoff Woodward, Oliver Lyttleton, David M Evans, Wolf Reik, Yu-Lee Paul, Gabriella Ficz, Susan E Ozanne, Anil Wipat, Keith Flanagan, Allyson Lister, Bastiaan T Heijmans, Susan M Ring, George Davey Smith; Data Resource Profile: Accessible Resource for Integrated Epigenomic Studies (ARIES), International Journal of Epidemiology, Volume 44, Issue 4, 1 August 2015, Pages 1181–1190, https://doi.org/10.1093/ije/dyv072

### GENR

Generation R is a population-based birth cohort aiming to identify early environmental and genetic determinants of development and health. All parents gave informed consent for their children's participation. The Generation R Study is conducted in accordance with the World Medical Association Declaration of Helsinki and study protocols have been approved by the Medical Ethics Committee of the Erasmus Medical Center, Rotterdam.

www.generationr.nl

Kooijman, M. N., Kruithof, C. J., van Duijn, C. M., Duijts, L., Franco, O. H., van IJzendoorn, M. H., ... & Moll, H. A. (2016). The Generation R Study: design and cohort update 2017. European journal of epidemiology, 31(12), 1243-1264.

### GLAKU

The adolescents of the Glaku (Glycyrrhizin in Licorice) cohort came from an urban community-based cohort comprising 1049 infants born between March and November 1998 in Helsinki, Finland (Strandberg et al., 2001). In 2009–2011, initial cohort members who had given permission to be contacted and whose addresses were traceable (N = 920, 87.7% of the original cohort in 1998) were invited to a follow-up, of which 692 (75.2%) could be contacted by phone (mothers of the adolescents). Of them, 451 (65.2% of those who could be contacted by phone, 49% of the invited) participated in a follow-up at a mean age of 12.3 years (SD = 0.5, range 11.0–13.2 years).

blogs.helsinki.fi/depsy-group/research/

Strandberg TE, Järvenpää AL, Vanhanen H, McKeigue PM. Birth outcome in relation to licorice consumption during pregnancy. Am J Epidemiol. 2001;153(11):1085-1088. doi:10.1093/aje/153.11.1085

### HELIX

The HELIX study represents a collaborative project across six established and ongoing longitudinal population-based birth cohort studies in Europe: the Born in Bradford (BiB) study in the UK, the Étude des Déterminants pré et postnatals du développement et de la santé de l'Enfant (EDEN) study in France, the INfancia y Medio Ambiente (INMA) cohort in Spain, the Kaunas cohort (KANC) in Lithuania, the Norwegian Mother, Father and Child Cohort Study (MoBa) and the RHEA Mother Child Cohort study in Crete, Greece. The HELIX project aims to measure and describe multiple environmental exposures from the different exposome domains during early life (pregnancy and childhood) and associate these with omics markers and child health outcomes.

www.projecthelix.eu

Maitre L, de Bont J, Casas M, Robinson O, Aasvang GM, Agier L, et al. 2018. Human Early Life Exposome (HELIX) study: a European population-based exposome cohort. BMJ Open 8:e021311; doi:10.1136/bmjopen-2017-021311.

### INMA

The INMA—INfancia y Medio Ambiente—(Environment and Childhood) Project is a network of birth cohorts in Spain that aim to study the role of environmental pollutants in air, water and diet during pregnancy and early childhood in relation to child growth and development (http://www.proyectoinma.org/) (Guxens et al. 2012). The study has been approved by Ethical

Committee of each participating centre and written consent was obtained from participating parents. Data for this study comes from INMA Sabadell subcohort.

www.proyectoinma.org

Guxens M, Ballester F, Espada M, et al. 2012. "Cohort Profile: the INMA--INfancia y Medio Ambiente--(Environment and Childhood) Project". International Journal of Epidemiology 41:930-40.

### NEST

Nest is a birth cohort recruited in six prenatal clinics who intended to use either Duke University or Durham Regional hospitals for their obstetrics care. The overarching goal was to identify early exposures associated with epigenomic alterations and phenotypic outcomes during childhood. All pregnant women were consented for their children's participation, and the protocol was approved by Duke University Medical Center Institutional Review Board.

Hoyo C, Murtha A, Schildkraut JM, Forman M, Overcash F, Jirtle RL, Kurtzberg J, Demark-Wahnefried W and Murphy, SK. Maternal folic acid use and offspring methylation profile in differentially methylated region (DMRs) regulating IGF2 imprinting. Epigenetics, 2011; 6(7):928-36. PMCID: PMC3154433.

Liu Y, Murphy SK, Murtha AP, Fuemmeler BF, Schildkraut JM, Huang Z, Overcash F, Kurtzberg J, Jirtle RL, Iversen ES, Forman MR, and Hoyo C. Depression in pregnancy, infant birth weight and DNA methylation of imprinted regulatory elements. Epigenetics. 2012; 7(7). PMCID: PMC3414394

### PREDO

The Prediction and Prevention of Preeclampsia and Intrauterine Growth Restriction (PREDO) is a prospective birth cohort study of Finnish women who were pregnant between 2005 and 2010 and their children. The PREDO study cohort was set up to identify novel risk factors and biomarkers in pregnant women associated with the development of preeclampsia and intrauterine growth restriction (IUGR), to (a) identify effective methods for prediction and prevention of preeclampsia in at-risk women, and (b) determine the association between exposure to preeclampsia, IUGR, or their risk factors and child developmental/health outcomes. Women with a singleton, intrauterine pregnancy who visited antenatal clinics at ten study hospitals in Finland for their first ultrasound screening at 12+0-13+6 weeks+days of gestation were recruited in the PREDO study. Two groups of pregnant women were enrolled: first, pregnant women with a known clinical risk factor status for preeclampsia and IUGR, and second, pregnant women who volunteered to participate regardless of their risk factor status for preeclampsia and IUGR. The sample with a known risk factor status comprises 1,079 pregnant women who gave live birth (969 of these women had at least one and 110 had none of the known risk factors for preeclampsia and IUGR). The community-based sample comprises 3,698 pregnant women who gave live birth. The sample with a known risk factor status visited antenatal clinics up to four times during pregnancy and both samples filled in bi-weekly self-reports. The post-delivery follow-up has taken place at approximately 2 weeks, 6 months, and 3.5 years after the delivery. The most recent follow-up started in 2016 and is ongoing. The study protocol was approved by the Ethics Committee of Obstetrics and Gynaecology, and Women, Children and Psychiatry of the Helsinki and Uusimaa Hospital District and by the participating hospitals. All participants provided written informed consent. Consent of participating children were provided by parent(s)/guardian(s).

In the high-risk sample, cord blood samples were collected according to standard procedures. DNA was extracted at the National Institute for Health and Welfare, Helsinki, Finland and and the Finnish Institute of Molecular Medicine, University of Helsinki, Finland and methylation analyses were performed at the Max Planck Institute in Munich, Germany.

blogs.helsinki.fi/depsy-group/research/
