## Supplementary Information 2 for "Association between DNA methylation and ADHD symptoms from birth to school age: A prospective meta-analysis"

#### Functional and molecular enrichment analysis

### Functional analysis

There are 249 probes showing suggestive evidence of association (P<1E-05) annotated to 182 unique genes.

Several GO-terms and KEGG pathways were nominally enriched among ADHD-suggestive CpGs. Three of them are of interest: The nicotinate and nicotinamide metabolism pathway (KEGG) (P-value=0.009, FDR=0.899), the riboflavin metabolism pathway (KEGG) (P-value=0.021, FDR=0.896) and the retrograde endocannabinoid signaling pathway (KEGG) (P-value=0.022, FDR=0.896).

Nicotinate (niacin) and nicotinamide are precursors of the coenzymes nicotinamide-adenine dinucleotide (NAD+) and nicotinamide-adenine dinucleotide phosphate (NADP+). These coenzymes, NAD+ and NADP+, are crucial for many metabolic pathways including glycolysis, TCA cycle, pentose phosphate cycle, fatty acid biosynthesis. When NAD+ and NADP+ are interchanged in a reaction with their reduced forms, NADH and NADPH respectively, they are important cofactors in several hundred redox reactions (Magni et al. 2004). Mechanisms related to fatty acid oxidation have been previously related to ADHD (Walton et al. 2017; Wilmot et al. 2016).

Riboflavin (vitamin B2, E101) is an essential component for the cofactors FAD (flavin-adenine dinucleotide) and FMN (flavin mononucleotide). Together with NAD+ and NADP+, FAD and FMN are important hydrogen carriers and take part in more than 100 redox reactions involved in energy metabolism (Rivlin 1970). Low levels of vitamin B2 were associated with ADHD diagnosis in adults (Landaas et al. 2016).

Endocannabinoids modulate synaptic function. By activating cannabinoid receptors expressed in the central nervous system, these lipid messengers can regulate several neural functions and behaviors. As experimental tools advance, the repertoire of known endocannabinoid-mediated effects at the synapse, and their underlying mechanism, continues to expand. Retrograde signaling is the principal mode by which endocannabinoids mediate short- and long-term forms of plasticity at both excitatory and inhibitory synapses (Castillo et al. 2012).

**GO collection:** Seven-hundred fifteen pathways were nominally associated with suggestive probes but none remained significant following FDR-correction. Full results in S1 (Supp_PATHWAYS_EWAS_ADHD_20181128.xls).

| | Term | Ont | N | DE | P.DE | FDR |
|---|---|---|---|---|---|---|
| **GO:0010256** | endomembrane system organization | BP | 413 | 14 | 5,32E-05 | 0,63202644 |
| **GO:0030953** | astral microtubule organization | BP | 7 | 3 | 5,67E-05 | 0,63202644 |
| **GO:0001968** | fibronectin binding | MF | 26 | 4 | 9,65E-05 | 0,68993315 |
| **GO:0006999** | nuclear pore organization | BP | 15 | 3 | 0,000258202 | 0,68993315 |
| **GO:0018995** | host | CC | 68 | 5 | 0,000299251 | 0,68993315 |
| **GO:0043657** | host cell | CC | 68 | 5 | 0,000299251 | 0,68993315 |
| **GO:0022411** | cellular component disassembly | BP | 549 | 15 | 0,000335307 | 0,68993315 |
| **GO:0044446** | intracellular organelle part | CC | 8568 | 105 | 0,000367984 | 0,68993315 |
| **GO:0044422** | organelle part | CC | 8793 | 107 | 0,000408591 | 0,68993315 |
| **GO:0044215** | other organism | CC | 74 | 5 | 0,000409846 | 0,68993315 |

**KEGG collection:** Seven pathways were nominally associated with suggestive probes but none remained significant following FDR-correction. Full results in S2 (Supp_PATHWAYS_EWAS_ADHD_20181128.xls).

| | **Pathway** | **N** | **DE** | **P.DE** | **FDR** |
|---|---|---|---|---|---|
| **path:hsa00770** | Pantothenate and CoA biosynthesis | 19 | 6 | 0,00678533 | 0,89591107 |
| **path:hsa00760** | Nicotinate and nicotinamide metabolism | 29 | 7 | 0,008778 | 0,89591107 |
| **path:hsa04621** | NOD-like receptor signaling pathway | 153 | 21 | 0,00987479 | 0,89591107 |
| **path:hsa04130** | SNARE interactions in vesicular transport | 33 | 7 | 0,01436319 | 0,89591107 |
| **path:hsa00740** | Riboflavin metabolism | 8 | 3 | 0,02099477 | 0,89591107 |
| **path:hsa04723** | Retrograde endocannabinoid signaling | 133 | 21 | 0,02190592 | 0,89591107 |
| **path:hsa05161** | Hepatitis B | 131 | 21 | 0,02246899 | 0,89591107 |
| **path:hsa05206** | MicroRNAs in cancer | 279 | 34 | 0,02372962 | 0,89591107 |
| **path:hsa04115** | p53 signaling pathway | 72 | 13 | 0,02443394 | 0,89591107 |
| **path:hsa00780** | Biotin metabolism | 3 | 2 | 0,02730377 | 0,90102448 |

### Molecular enrichment analysis

**Summary**

ADHD-FDR CpGs were enriched for 3'UTR and body regions and depleted for Tss1500, Tss200 and first exon regions. They were also enriched in open sea positions and N- and S-shelfs, and depleted in N- and S-shores and CpG islands. The same pattern was observed for hypomethylated FDR CpGs, and the inverse for hypermethylated FDR CpGs. Regarding blood 15-chormatine states, FDR hypomethylated CpGs showed enrichment for transcription (TxWk, Tx), enhancers (EnhG, Enh), and ZNF genes and repeats and quiescent positions, while depletion for transcription start sites (TssA, TssAFlnk, TxFlnk), bivalent (TssBiv, BivFlnk, EnhBiv) and repressor (ReprPC) positions. The pattern for hypermehylated FDR CpGs was inverted. Overall, this indicates that ADHD-associated hypomethylation tends to happen in enhancers, transcribed regions, and quiescent regions, while hypermethylation tends to happen in transcription start sites and bivalent state regions.
Similar enrichment/depletion patters were observed with suggestive CpGs.
See below the detailed findings from the molecular enrichment analysis.

**Genic vs. intergenic probes**

Suggestive CpGs, specially the hypomethylated, were enriched in intergenic regions.

| SUGGESTIVE | Genic (N=355420) | Intergenic (N=117397) |
|---|---|---|
| **Sig SUGG (N=249)** | 170 (0.05%) | 79 (0.075%) |
| **No sig SUGG (N=472568)** | 355250 (99.95%) | 117318 (99.93%) |

X-squared = 5.9862, df = 1, p-value = 0.01442

Odds Ratio: 0.8904

| | Genic (N=355420) | Intergenic (N=117397) |
|---|---|---|
| **Hyper SUGG (N=39)** | 28 | 11 |
| **Rest (N=472778)** | 355392 | 117386 |

X-squared = 0.091614, df = 1, p-value = 0.7621

Odds Ratio: 1.1893

| | Genic (N=355420) | Intergenic (N=117397) |
|---|---|---|
| **Hypo SUGG (N=210)** | 142 | 68 |
| **Rest (N= 472607)** | 355278 | 117329 |

X-squared = 6.0209, df = 1, **p-value = 0.01414**

Odds Ratio: 0.6896305

**Relative gene position**

Suggestive CpGs were enriched for 3'UTR regions and depleted for TSS200 and first exon regions. The same is observed for hypomethylated.

| | **Suggestive** | |
|---|---|---|
| | **Yes** | **No** |
| **Hyper** | 39 | 239939 |
| **Hypo** | 210 | 232629 |

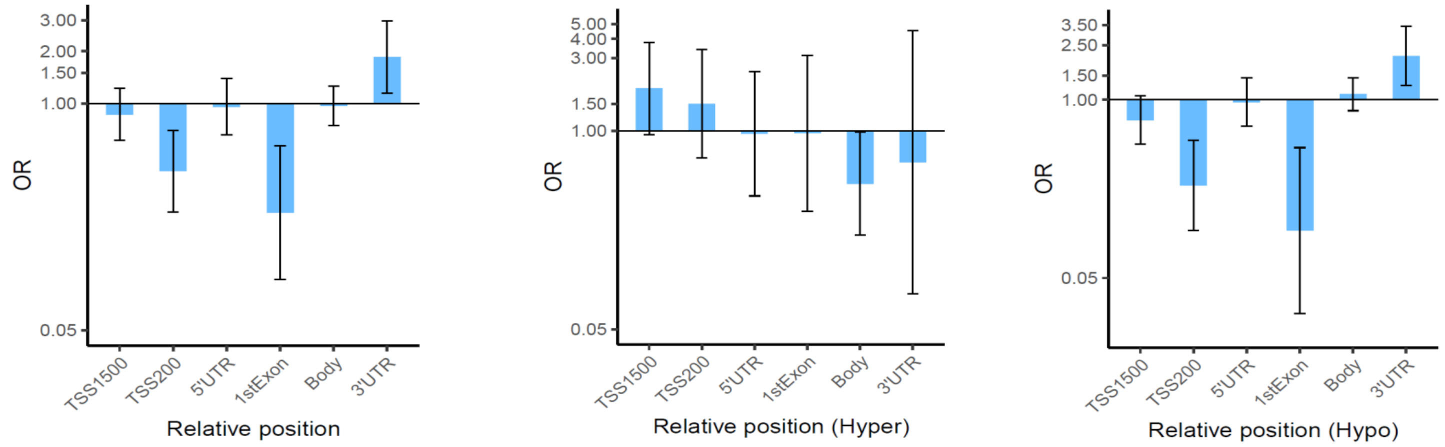


**Figure 1.** Bar plots depicting OR and 95% CIs for suggestive significant (left), hypermethylated suggestive (center) and hypomethylated suggestive probes (right) compared to no suggestive significant probes in relation to genic position.

**Relation to CpG Island**

Suggestive CpGs were enriched for open sea, north shelf and south shelf regions and depleted for south shore and islands. This pattern is manteined for hypomethylated CpGs.

| Suggestive | Island | North Shelf | North Shore | Open Sea | South Shelf | South Shore |
|---|---|---|---|---|---|---|
| No sig sugg | 145756 (30.8%) | 24110 (5.1%) | 61104 (12.9%) | 172193 (36.4%) | 21629 (4.6%) | 47776 (10.1%) |
| Sig sugg | 10 (4.0%) | 25 (10.0%) | 22 (8.8%) | 161 (64.7%) | 21 (8.4%) | 10 (4.0%) |

X-squared = 144.82, df = 5, p-value < 2.2e-16

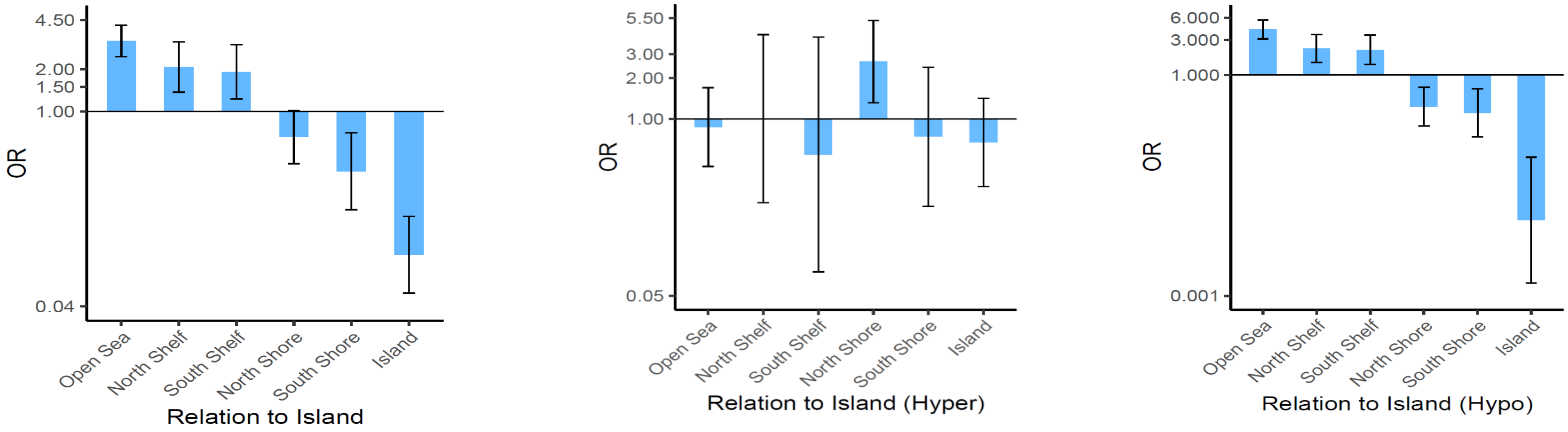


**Figure 2.** Bar plots depicting OR and 95% CIs for suggestive significant (left), hypermethylated suggestive (center) and hypomethylated suggestive CpGs (right) compared to no suggestive significant CpGs in relation to CpG island.

**Chromatin states**

Both suggestive CpGs and hypomethylated suggestive CpGs showed the same pattern: enrichment for transcription (Tx and TxWk) and quiescent positions and depletion for transcription start site positions (TSSA, TxFlnk, TxFlnk) and bivalent (EnhBiv) and repressor (ReprPC) positions. Overall, hypermethylated suggestive CpgS showed an opposite pattern of results compared to hypomethylated CpGs. The states are described in Page 2.

| | Suggestive | |
|---|---|---|
| | **Yes** | **No** |
| **Hyper** | 39 | 239939 |
| **Hypo** | 210 | 232629 |

-

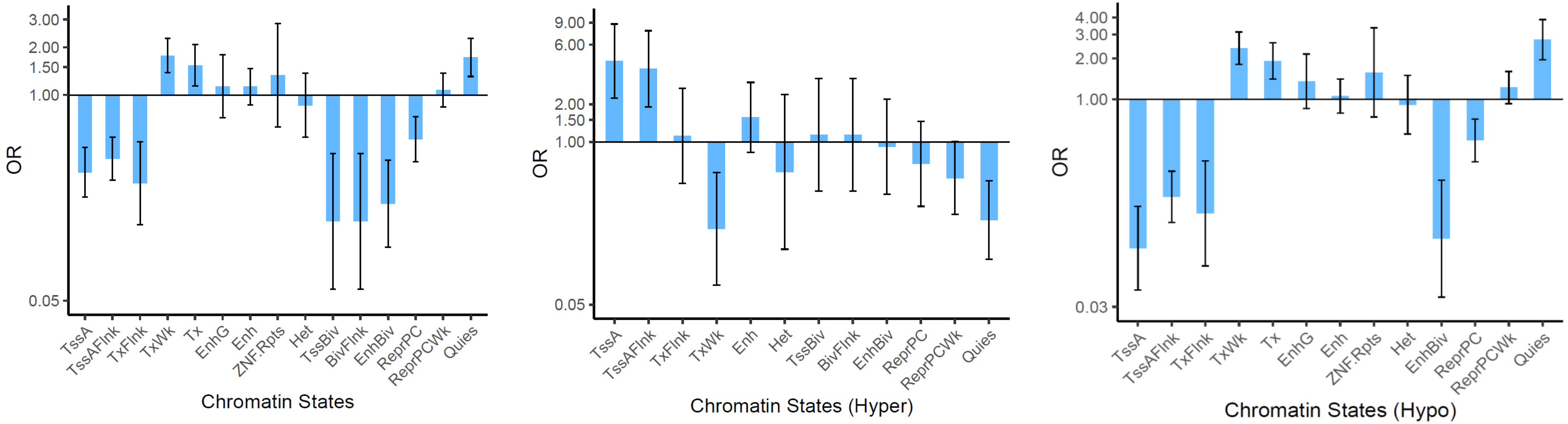


**Figure 3.** Bar plots depicting OR and 95% CIs for suggestive significant (left), hypermethylated suggestive (center) and hypomethylated suggestive probes (right) compared to no suggestive significant probes for the different chromatin states.
