## Supplementary Tables S2-S4 for "Association between DNA methylation and ADHD symptoms from birth to school age: A prospective meta-analysis"

*Table S2: Phenotype Characteristics*

| Study Name | Ancestry/Ethnicity | Instrument | EWAS | Assesment age (mean years) | Age (SD) | Total ADHD score (mean) | Total ADHD score (SD) | % Girls |
|---|---|---|---|---|---|---|---|---|
| ALSPAC | European | DAWBA | Birth and school-age | 7.5 | 0.9 | 0.5 | 0.9 | 52 |
| ALSPAC | European | DAWBA | Birth | 10.7 | 0.1 | 0.5 | 0.9 | 51 |
| ALSPAC | European | DAWBA | Birth | 13.9 | 0.1 | 0.5 | 0.9 | 50 |
| ALSPAC | European | DAWBA | Birth | 15.4 | 0.2 | 0.5 | 0.9 | 51 |
| GENR | European | CBCL 1.5-5 | Birth and school-age | 5.9 | 0.3 | 2.7 | 2.4 | 51 |
| GENR | European | Conners | Birth | 8.1 | 0.2 | 7.5 | 6.6 | 51 |
| GENR | European | CBCL 6-18 | Birth | 9.7 | 0.3 | 2.6 | 2.8 | 51 |
| GLAKU | European | CBCL | School age | 12.3 | 0.5 | 1.9 | 2.4 | 54.6 |
| HELIX | European | Conners | School-age | 8.0 | 1.6 | 7.8 | 6.6 | 44.7 |
| HELIX | Pakistani | Conners | School-age | 6.6 | 0.2 | 6.7 | 6.9 | 44.3 |
| INMA | European | Conners | Birth | 6.7 | 0.4 | 7.7 | 6.9 | 52 |
| INMA | European | CBCL 6-18 | Birth | 8.9 | 0.6 | 3.4 | 3.1 | 48 |
| NEST | Black | BASC | Birth | 4.8 | 0.5 | 15.8 | 5.4 | 46 |
| NEST | White | BASC | Birth | 5.0 | 0.6 | 15.6 | 6.5 | 49 |
| PREDO | European | Conners | Birth | 4.5 | 0.5 | 5.0 | 4.4 | 47.1 |

*Table S3: Twin heritability of genome-wide significant CpG sites*

| CpG | A | C | E |
|---|---|---|---|
| cg01271805 | 0.19 | 0.10 | 0.71 |
| cg09158638 | 0.00 | 0.06 | 0.94 |
| cg09762907 | 0.02 | 0.09 | 0.89 |
| cg11251614 | 0.00 | 0.07 | 0.93 |
| cg17876201 | 0.00 | 0.09 | 0.91 |
| cg21600027 | 0.00 | 0.04 | 0.96 |
| cg22997238 | 0.00 | 0.25 | 0.75 |
| cg24838839 | 0.15 | 0.00 | 0.85 |
| cg25520701 | 0.00 | 0.11 | 0.89 |

Based on Hannon E, Knox O, Sugden K, Burrage J, Wong CCY, Belsky DW *et al.* Characterizing genetic and environmental influences on variable DNA methylation using monozygotic and dizygotic twins. *PLoS Genet* 2018; **14**: 1–27.

*Table S4: Replication of Walton et al. EWAS (ADHD trajectories and cord blood methylation)*

| | | | | Discovery | | | | Replication | | | | |
|---|---|---|---|---|---|---|---|---|---|---|---|---|
| CpG | Gene | Chr | Position | $n_{studies}$ | n | B | p | $n_{studies}$ | n | B | SE | p |
| cg18587973 | CDADC1 | 13 | 49822535 | 1 | 817 | 0.17 | 1.2E-06 | 5 | 1755 | 3.54 | 1.55 | 0.03 |
| cg27469152 | EPX | 17 | 56282313 | 1 | 817 | -0.18 | 2.0E-07 | 5 | 1763 | -1.20 | 0.85 | 0.20 |
| cg16290904 | PEX2 | 8 | 77912348 | 1 | 817 | 0.17 | 7.4E-07 | 5 | 1761 | -2.99 | 2.60 | 0.23 |
| cg03905179 | MAFK | 7 | 1582588 | 1 | 817 | 0.17 | 1.3E-06 | 5 | 1756 | 0.70 | 1.67 | 0.46 |
| cg24843380 | ZNF454 | 5 | 178367827 | 1 | 817 | 0.17 | 1.6E-06 | 5 | 1762 | -1.73 | 2.05 | 0.47 |
| cg05653018 | ELF3 | 1 | 201979533 | 1 | 817 | 0.17 | 1.4E-06 | 5 | 1763 | 0.45 | 0.60 | 0.53 |
| cg15096815 | JUN | 1 | 59249838 | 1 | 817 | -0.18 | 3.5E-07 | 5 | 1763 | 0.55 | 1.08 | 0.68 |
| cg24481594 | SKI | 1 | 2190850 | 1 | 817 | -0.20 | 1.5E-08 | 5 | 1763 | -0.42 | 1.08 | 0.76 |
| cg26263766 | ZNF544 | 19 | 58739734 | 1 | 817 | 0.17 | 8.7E-07 | 5 | 1714 | -0.31 | 1.13 | 0.84 |
| cg01324543 | CCDC30 | 1 | 42999439 | 1 | 817 | -0.17 | 7.2E-07 | 5 | 1763 | 0.59 | 1.75 | 0.92 |
| cg13714586 | FBXW5 | 9 | 139838358 | 1 | 817 | 0.17 | 1.3E-06 | 5 | 1752 | 0.72 | 4.50 | 0.93 |
| cg09989037 | ST3GAL3 | 1 | 44300942 | 1 | 817 | -0.17 | 9.5E-07 | 5 | 1763 | -0.17 | 0.56 | 0.96 |
| cg22193912 | MAFG | 17 | 79881523 | 1 | 817 | 0.17 | 1.3E-06 | 5 | 1763 | 0.24 | 0.81 | 0.99 |

**Chr** Chromosome
**$n_{studies}$** Number of studies
**n** Number of participants
**B** Regression coefficient
SE Standard error

*Table S5: Replication of Wilmot et al. EWAS (case-control study in school-age)*

| | | | | Discovery | | | | Replication | | | | |
|---|---|---|---|---|---|---|---|---|---|---|---|---|
| CpG | Gene | Chr | Position | $n_{studies}$ | n | Diff. | p | $n_{studies}$ | n | B | SE | p |
| cg05180887 | MYT1L | 2 | 1817263 | 1 | 92 | -0.05 | 0.04 | 4 | 2080 | -0.18 | 0.20 | 0.43 |
| cg08479516 | VIPR2 | 7 | 158905536 | 1 | 92 | -0.03 | 0.03 | 4 | 1900 | 0.04 | 0.52 | 0.57 |
| cg06201514 | MYT1L | 2 | 1817409 | 1 | 92 | -0.08 | 0.02 | 5 | 2295 | -0.06 | 0.11 | 0.66 |
| cg13444538 | VIPR2 | 7 | 158905317 | 1 | 92 | -0.06 | 0.03 | 4 | 2080 | -0.06 | 0.23 | 0.74 |
| cg10075506 | MYT1L | 2 | 1817351 | 1 | 92 | -0.08 | 0.04 | 5 | 2295 | -0.02 | 0.12 | 0.88 |
| cg05554000 | VIPR2 | 7 | 158905015 | 1 | 92 | -0.05 | 0.02 | 5 | 2295 | -0.02 | 0.16 | 0.92 |

**Chr** Chromosome
**$n_{studies}$** Number of studies
**n** Number of participants
**Dff.** Difference in methylation between cases and controls (Negative values indicate hypomethylation in controls)
**B** Regression coefficient
**SE** Standard error
